# Structural basis for the dimerization-dependent CRISPR-Cas12f nuclease

**DOI:** 10.1101/2020.12.22.424058

**Authors:** Renjian Xiao, Zhuang Li, Shukun Wang, Ruijie Han, Leifu Chang

## Abstract

Cas12f, also known as Cas14, is an exceptionally small type V-F CRISPR-Cas nuclease that is roughly half the size of comparable nucleases of this type. To reveal the mechanisms underlying substrate recognition and cleavage, we determined the cryo-EM structures of the Cas12f-sgRNA-target DNA and Cas12f-sgRNA complexes at 3.1 Å and 3.9 Å, respectively. An asymmetric Cas12f dimer is bound to one sgRNA for recognition and cleavage of dsDNA substrate with a T-rich PAM sequence. Despite its dimerization, Cas12f adopts a conserved activation mechanism among the type V nucleases which requires coordinated conformational changes induced by the formation of the crRNA-target DNA heteroduplex, including the close-to-open transition in the lid motif of the RuvC domain. Only one RuvC domain in the Cas12f dimer is activated by substrate recognition, and the substrate bound to the activated RuvC domain is captured in the structure. Structure-assisted truncated sgRNA, which is less than half the length of the original sgRNA, is still active for target DNA cleavage. Our results expand our understanding of the diverse type V CRISPR-Cas nucleases and facilitate potential genome editing applications using the miniature Cas12f.

## INTRODUCTION

Clustered regularly interspaced short palindromic repeats (CRISPR) and CRISPR-associated proteins (Cas) systems are the adaptive immune systems in bacteria and archaea against infection from mobile genetic elements (MGEs) ^1-4^. In Class 2 CRISPR-Cas systems, a single effector nuclease incorporates with guide RNAs (gRNAs) to recognize target DNA with complementary sequences. Class 2 systems are further divided into three types: type-II exemplified by Cas9 nucleases, type-V featured by Cas12 nucleases, and type VI epitomized by Cas13 nucleases ^5^. Type-V systems have the most subtypes discovered to date, including Cas12a-k ^5,6^. Most Cas12 nucleases target double-stranded DNA for cleavage, with the exception of Cas12g which targets RNA substrates ^6^ and Cas12k which is inactive for substrate cleavage ^7^.

Cas12f, also known as Cas14, is the smallest class 2 CRISPR-Cas effector reported to date with a length between ∼400-700 amino acids ^8^. Cas12f proteins were identified almost exclusively within a superphylum of symbiotic archaea, DPANN ^8^. Initially found to be specific for ssDNA, Cas12f was recently reported to also recognize dsDNA with 5’ T-rich protospacer adjacent motifs (PAMs) ^9^. Cas12f associates with a crRNA and a tracrRNA, which can be fused into a single guide RNA (sgRNA), to target substrate DNA. Cas12f is a Mg^2+^-dependent endonuclease that functions best in low salt concentrations and at ∼46°C ^9^. Similar to other Cas12 nucleases, Cas12f is capable of cleaving non-specific ssDNA in *trans* after binding complementary target DNA, thus enabling its development for nucleic acid detection ^8^.

Type-V effectors employ multiple domains distributed in a recognition lobe (REC) and a nuclease lobe (NUC) for substrate recognition and cleavage. The REC lobe is responsible for substrate recognition, whereas the NUC lobe contains a nuclease site located within the RuvC domain. Most Type-V effectors contain about 1000 amino acids, such as Cas12a (1226-1307 a.a.) ^10-20^, Cas12b (1108-1129 a.a.) ^21-23^, Cas12e (986 a.a.) ^24^, and Cas12i (1093 a.a.) ^25^. However, it is not known how a miniature Cas12f, which is only about half the size of other Cas12 nucleases, completes all functional requirements for target recognition and cleavage. By determining the atomic structures of Cas12f-sgRNA in the presence and absence of target dsDNA, we show here that two copies of Cas12f are required for substrate recognition and cleavage.

## MATERIALS AND METHODS

### Protein expression and purification

The plasmid encoding full-length Cas12f (UnCas12f1) was purchased from Addgene #112500 with an N-terminal 10xHis-MBP-tag. The plasmid was transformed into *E. coli* BL21(DE3) cells and grown to OD_600_ =0.5 in Terrific Broth (TB). Protein overexpression was induced by adding 0.5 mM IPTG followed by incubation at 18°C overnight. The cells were collected and then resuspended in buffer A containing 25 mM Tris-HCl (pH 7.6), 1 M NaCl, 5% glycerol, 1 mM PMSF, and 5 mM β-mercaptoethanol, and disrupted by sonication. Cell lysate was clarified by centrifugation. The supernatant was loaded onto Ni-NTA resin, washed with buffer B containing 25 mM Tris-HCl (pH 7.6), 1 M NaCl, 30 mM imidazole, and 5 mM β-mercaptoethanol, and the Cas12f protein was eluted by buffer B supplemented with 250 mM imidazole. The His-MBP-tag was removed by overnight incubation with TEV protease at 4°C. The target protein was exchanged into buffer C containing 25 mM Tris-HCl (pH 7.6), 500 mM NaCl, 2 mM DTT, and 5% glycerol, loaded onto a HiTrap SP HP column (GE Healthcare), and eluted with a linear NaCl gradient (0.1 to 2M) followed by size exclusion chromatography over a Superdex 200 (GE Healthcare) in buffer D containing 25 mM Tris-HCl (pH 7.6), 150 mM NaCl, 2 mM DTT, and 1 mM MgCl_2_. Fractions were concentrated and stored at −80°C.

To assemble the Cas12f-sgRNA binary complex, Cas12f proteins were incubated with sgRNA **(Table S1)** at a ratio of 1:1.2 at 37°C for 30 min in buffer D. To reconstitute the Cas12f-sgRNA-target DNA complex, Cas12f D510A mutant proteins were incubated with guide RNA at 37°C for 30 min followed by adding the target DNA **(Table S1)** synthesized from IDT at a ratio of 1:1.2:1.3. After 30 min, the reaction mixture was subjected to SEC over a Superdex 200 column (GE Healthcare) equilibrated with buffer D for further purification.

### sgRNA preparation

sgRNAs were produced by *in vitro* transcription using the HiScribe T7 High Yield RNA synthesis kit (NEB) with PCR amplified gBlocks (IDT) as templates. sgRNAs were purified over a Resource-Q column (GE Healthcare) and eluted with a linear NaCl gradient (50 mM-1000 mM) in 25 mM Tris–HCl (pH 8.0). The eluted sgRNAs were concentrated and stored at −80°C

### Mutagenesis

Single amino acid mutations were introduced by the QuikChange site-directed mutagenesis method. Mutations with multiple amino acids were introduced by ligating inverse PCR-amplified backbone with mutations bearing DNA oligonucleotides via the In-Fusion Cloning Kit (ClonTech). All mutants were confirmed by Sanger sequencing.

### *In vitro* DNA cleavage assay

Target DNA containing the 5′-TTTA-3′ PAM was ordered from IDT and cloned into a pET28-MHL vector using the In-Fusion Cloning Kit (ClonTech). Plasmids were linearized before usage. Cas12f proteins (200 nM) were mixed with guide RNA at a ratio of 1:1.1 at 37°C for 30 min in cleavage buffer containing 2.5 mM Tris-HCl (pH 7.6), 50 mM NaCl, 10 mM MgCl_2_, and 0.5 mM DTT, and then linearized plasmids (5 nM) were added. The reactions were quenched by adding EDTA and proteinase K (Thermo Fisher Scientific) after 45 min. The cleavage products were resolved on 0.7% agarose gels and visualized by ethidium bromide staining.

### Electron Microscopy

Aliquots of 4 μL Cas12f-sgRNA binary complex (1 mg/mL) and Cas12f-sgRNA-dsDNA ternary complex (1 mg/mL) were applied to glow-discharged UltrAuFoil holey gold grids (R1.2/1.3, 300 mesh). The grids were blotted for 2 s and plunged into liquid ethane using a Vitrobot Mark IV. Cryo-EM data were collected with a Titan Krios microscope (FEI) operated at 300 kV and images were collected using Leginon ^26^ at a nominal magnification of 81,000x (resulting in a calibrated physical pixel size of 1.05 Å/pixel) with a defocus range of −0.8 – −2.0 μm. The images were recorded on a K3 electron direct detector in super-resolution mode at the end of a GIF-Quantum energy filter operated with a slit width of 20 eV. A dose rate of 20 electrons per pixel per second and an exposure time of 3.12 seconds were used, generating 40 movie frames with a total dose of ∼ 54 electrons per Å^2^. Statistics for cryo-EM data are listed in **Table 1**.

**Table 1.** Cryo-EM data collection, refinement and validation statistics.

|  | Cas12f-sgRNA-target DNA | Cas12f-sgRNA |
| --- | --- | --- |
| **Data collection and processing** |  |  |
| Magnification | 81,000 | 81,000 |
| Voltage (kV) | 300 | 300 |
| Electron exposure (e<sup>-</sup>/Å<sup>2</sup>) | 54 | 54 |
| Defocus range (-μm) | 0.8 – 2.0 | 0.8 – 2.0 |
| Pixel size (Å) | 1.05 | 1.05 |
| Symmetry imposed | C1 | C1 |
| Initial particle images (no.) | 3,284,618 | 1,846,279 |
| Final particle images (no.) | 384,132 | 154,090 |
| Map resolution (Å) | 3.1 | 3.9 |
| FSC threshold | 0.143 | 0.143 |
| Map resolution range (Å) | 2.8 – 4.0 | 3.7 – 4.9 |
| **Refinement** |  |  |
| Initial model used | None | PBD:7L49 |
| Model resolution (Å) | 3.1 | 3.9 |
| FSC threshold | 0.5 | 0.5 |
| Model resolution range (Å) | 2.8 – 4.0 | 3.7 – 4.9 |
| Map sharpening *B* factor (Å<sup>2</sup>) | -97 | -117 |
| Model composition |  |  |
| Non-hydrogen atoms | 12330 | 10889 |
| Protein residues | 1041 | 1041 |
| Nucleotides | 187 | 116 |
| Ligands | 4 (Zn) | 4 (Zn) |
| *B* factors (Å<sup>2</sup>) |  |  |
| Protein | 86.88 | 440.91 |
| Nucleotide | 164.05 | 398.21 |
| Ligands | 95.33 | 418.46 |
| R.m.s. deviations |  |  |
| Bond lengths (Å) | 0.003 | 0.003 |
| Bond angles (°) | 0.758 | 0.756 |
| **Validation** |  |  |
| MolProbity score | 2.16 | 2.26 |
| Clashscore | 13.85 | 15.34 |
| Poor rotamers (%) | 0.33 | 0.11 |
| Ramachandran plot |  |  |
| Favored (%) | 91.42 | 89.20 |
| Allowed (%) | 8.29 | 10.22 |
| Disallowed (%) | 0.29 | 0.58 |

### Image Processing

The movie frames were imported to RELION-3 ^27^. Movie frames were aligned using MotionCor2 ^28^ with a binning factor of 2. Contrast transfer function (CTF) parameters were estimated using Gctf ^29^. A few thousand particles were auto-picked without template to generate 2D averages for subsequent template-based auto-picking. The auto-picked and extracted particle datasets were split into batches for 2D classifications, which were used to exclude false and bad particles that fell into 2D averages with poor features. Particles from different views were used to generate an initial model in cryoSPARC ^30^. 3D classification was further performed to distinguished different compositional/conformational heterogeneity. The homogeneous dataset was used for final 3D refinement with C1 symmetry.

For the Cas12f-sgRNA binary complex dataset, 1,846,279 particles were auto-picked and extracted from 1,391 dose weighted micrographs. 448,190 particles were selected from 2D classification and used for 3D classification. 154,190 particles were selected from 3D classification and used for final 3D refinement.

For the Cas12f-sgRNA-dsDNA ternary complex dataset, 3,284,618 particles were auto-picked and extracted from 2,450 dose weighted micrographs. 992,872 particles were selected from 2D classification and used for 3D classification. 384,132 particles were selected from 3D classification and used for final 3D refinement. Focused refinement around the Nuc domain was further performed to improve the local map quality.

Cryo-EM image processing is summarized in **Table 1**.

### Model building, refinement, and validation

*De novo* model building of the Cas12f-sgRNA-target DNA structure was performed manually in COOT ^31^ guided by secondary structure predictions from PSIPRED ^32^. Refinement of the structure models against corresponding maps were performed using the *phenix*.*real_space_refine* tool in Phenix ^33^. For the Cas12f-sgRNA complex, the structure model of the Cas12f-sgRNA-target-DNA complex was fitted into the cryo-EM map, and each domain was manually adjusted in COOT. The resultant model was refined against the corresponding cryo-EM map using the *phenix*.*real_space_refine* tool in Phenix. 3D FSC analysis for the presented maps were performed using the Remote 3DFSC Processing Server (https://3dfsc.salk.edu/upload/) ^34^.

### Structural visualization

Figures were generated using PyMOL and UCSF Chimera ^35^.

## RESULTS

### Overall structure of Cas12f-sgRNA-target DNA

We assembled a Cas12f-sgRNA-target DNA ternary complex by incubating an inactive Un1Cas12f1^D510A^ (529 amino acids or a.a., 61.5 kDa)^9^, a sgRNA (222 nucleotides), and a target dsDNA with a TTTA PAM sequence (60 base pairs) **(Fig. S1A)**. Using cryo-EM, we determined the structure of this complex at 3.1 Å resolution **(Figs. 1A and S1B-G, and Table 1)**. The resultant map allowed us to build the atomic model of the whole complex **(Fig. S2)**, except three residues at the N-terminus, four residues at the C-terminus, and flexible regions in the sgRNA and target DNA to be discussed below. The most astonishing feature of the structure is the presence of two copies of Cas12f in the complex (named Cas12f.1 and Cas12f.2) **(Fig. 1A,B)**, in contrast to all previous determined structures of other class 2 effectors.

**Figure 1.**
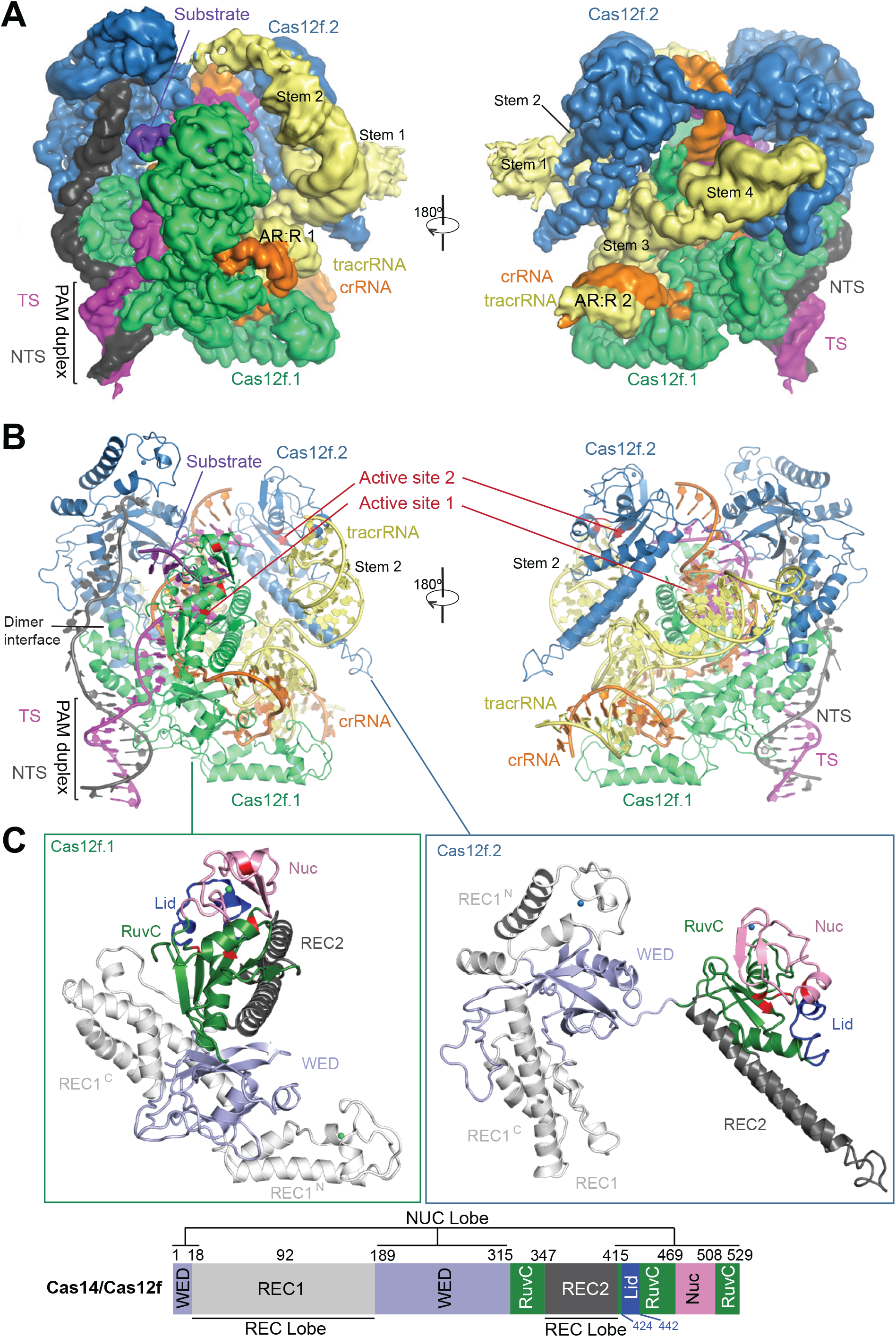
Overall structure of the Cas12f-sgRNA-target DNA complex. **(A)** Cryo-EM map of the Cas12f-sgRNA-target DNA complex at 3.1 Å in two views with each subunit color coded (Cas12f.1 in green, Cas12f.2 in sky blue, crRNA in orange, tracrRNA in pale yellow, target and non-target strands in magenta and black, respectively). **(B)** Atomic model of the Cas12f-sgRNA-target DNA complex shown in cartoon. **(C)** Structure of Cas12f.1 and Cas12f.2 with each domain color-coded (REC1 in white, REC2 in black, WED in light blue, RuvC in forest, and Nuc in pink). The catalytic residues and lid motif in the RuvC domain are colored in red and blue, respectively. Schematic of domain organization of Cas12f is shown at the bottom.

The overall structure of the Cas12f-sgRNA-target DNA ternary complex is consistent with a recent study ^36^ that was published during the preparation of this paper. Despite its small size, Cas12f contains all the conventional domains of Cas12 proteins, compared with other known Cas12 nuclease structures **(Fig. S3)**. Cas12f monomers consist of REC1 and WED domains in the N-terminal half and the RuvC, REC2 (included as part of RuvC in ^36^), and Nuc (the target nucleic acid-binding or TNB domain in ^36^) domains in the C-terminal half **(Fig. 1C)**. The closest match to Cas12f is Cas12g with 767 amino acids with both Cas12f and Cas12g being classified into branch 3 of type V nucleases based on phylogenetic analysis ^6^. The biggest difference is the REC1 domain, which can be further divided into two subdomains: REC1^N^ (referred to as a zinc finger or ZF domain in ^36^) and REC1^C^. REC1^N^ contains two anti-parallel helices connected by a CCCH zinc finger motif while REC1^C^ is composed of a three anti-parallel helical bundle, which is the primary dimerization interface of Cas12f **(Fig. 1C)**.

### Structure of sgRNA

The sgRNA of Cas12f contains a 140-nt tracrRNA at the 5′ end and a 37-nt crRNA at the 3′ end (17-nt repeat-derived and 20-nt spacer-derived sequences), connected by a linker **(Fig. 2A,B)**. Four stem-loop structures (Stems 1-4) are present in the tracrRNA **(Figs. 1A and 2A,B)**. Stem 1 (1-21) contains seven base pairs and is solvent exposed but lacks direct interactions with the Cas12f subunits. Deletion of the Stem 1 (ΔStem 1) shows comparable activity to the full-length tracrRNA in substrate cleavage assays **(Fig. 2C)**. Stem 2 (22-69) is a long duplex primarily interacting with Cas12f.2 that connects the N-terminal and C-terminal halves of Cas12f.2 **(Fig. 1A)**. The 10-bp duplex (23-33 and 59-69) bound to the C-terminal half of Cas12f.2 is structurally ordered while the rest (34-58) is curved and flexible due to disturbance of the Watson-Crick base pairing in the duplex **(Figs.1A and 2B)**. Partial deletion of Stem 2 (ΔStem 2^34-58^) or both Stems 1 and 2 (ΔStems 1&2) results in reduced activity in substrate cleavage assays **(Fig. 2C)**, indicating that Stem 2 is required for optimal activity. The 5-bp Stem 3 (72-88) is located in the center of the sgRNA structure and contributes a loop (78-83) that forms the anti-repeat:repeat duplex 1 (AR:R 1) with the repeat-derived region of crRNA, critical for correct positioning of the spacer-derived guide **(Fig. 2A,B)**. Following Stem 3 is a long duplex Stem 4 (94-127) that lies between the two copies of Cas12f and establishes extensive interactions with both of them. Consequently, replacement of Stem 4 with a UUUU linker (ΔStem 4) significantly reduces substrate cleavage activity **(Fig. 2C)**. The 3’ end of the tracrRNA (132-140) establishes the second duplex with the repeat-derived region of crRNA, AR:R 2 duplex. Deletion of AR:R 2 duplex (ΔAR:R 2) shows moderate reduction in substrate cleavage activity. All together with the exception of Stem 1, the stem-loop structures in tracrRNA play a role in Cas12f activity. However, none of the deletion mutations completely abolish the complex’s activity. Notably, deletion of Stems 1 and 2, and AR:R 2 (ΔStems 1&2 & AR:R 2), reducing the sgRNA from 222-nt to 90-nt, still shows considerable substrate cleavage activity **(Fig. 2C)**. These results lay the foundation for designing smaller and simpler guide RNAs for potential application of Cas12f in genome editing.

**Figure 2.**
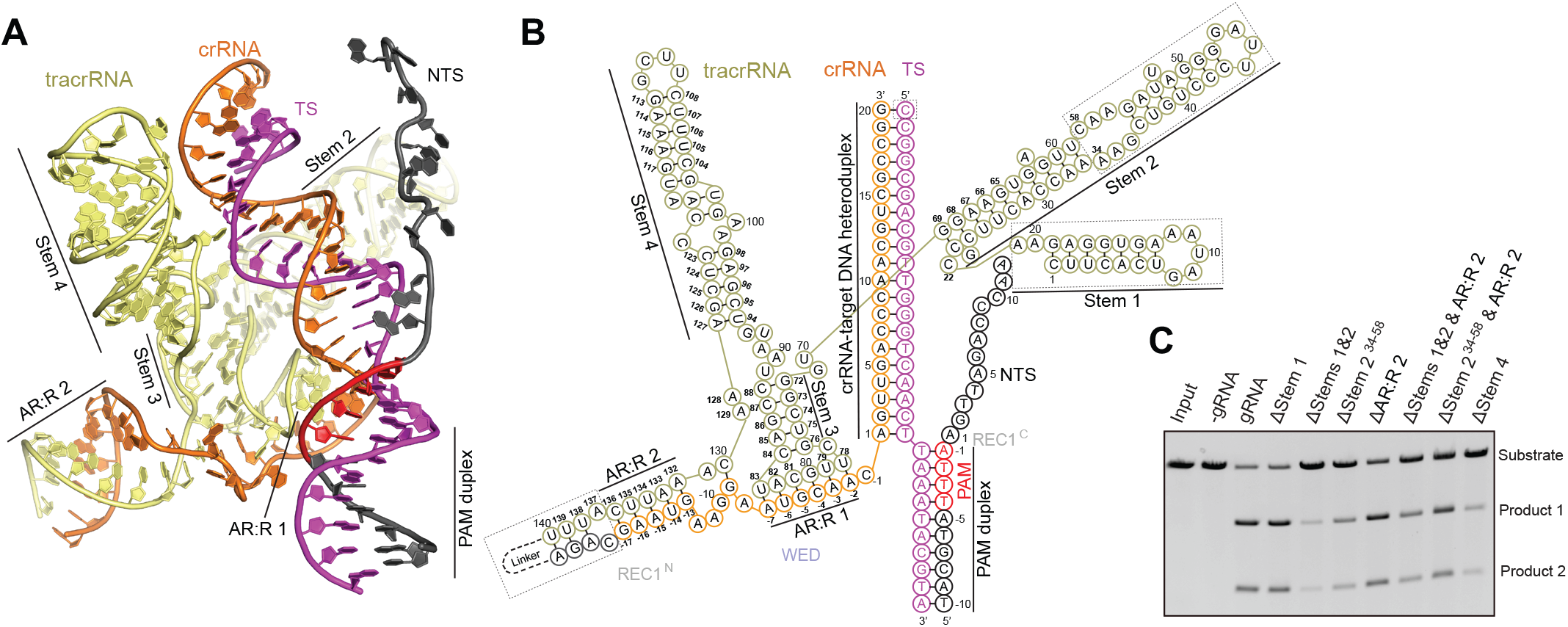
Overall structure of sgRNA. **(A)** Structure of the sgRNA and target DNA in the Cas12f-sgRNA-target DNA complex in cartoon presentation. **(B)** Schematic of the sgRNA and target DNA. Structurally disordered regions are indicated by dashed boxes. **(C)** DNA substrate cleavage assay using full-length sgRNA and sgRNA with various deletion mutants. The results shown are representative of three experiments.

### Dimerization

REC1^C^ forms a dimerization interface between two Cas12f monomers through hydrophobic interactions **(Figs. 1B and 3A)**. Specifically, five hydrophobic amino acids (I118, Y121, Y122, Y126, and L182) from each monomer establish a hydrophobic patch that associates the two monomers. Mutation of any of those residues to glycine reduces the cleavage activity of Cas12f, with Y121G, I126G, and L182G exhibiting significant effects **(Fig. 3B)**. Furthermore, mutations of two residues (Y121 and Y122) or four residues (I118, Y121, Y122, and Y126) to either glycine or glutamic acid completely abolish the cleavage activity **(Fig. 3B)**. These results suggest dimerization is essential for substrate cleavage by Cas12f. In addition to REC1^C^, the REC2 domains form a second contact between Cas12f.1 and Cas12f.2 through electrostatic interactions and their contacts with the Stem 4 of tracrRNA **(Fig. 3C)**. Except the dimerization interfaces, both Cas12f molecules establish extensive interactions with one copy of sgRNA, suggesting that sgRNA plays an important role for coordinating the two Cas12f molecules within the complex ^36^.

**Figure 3.**
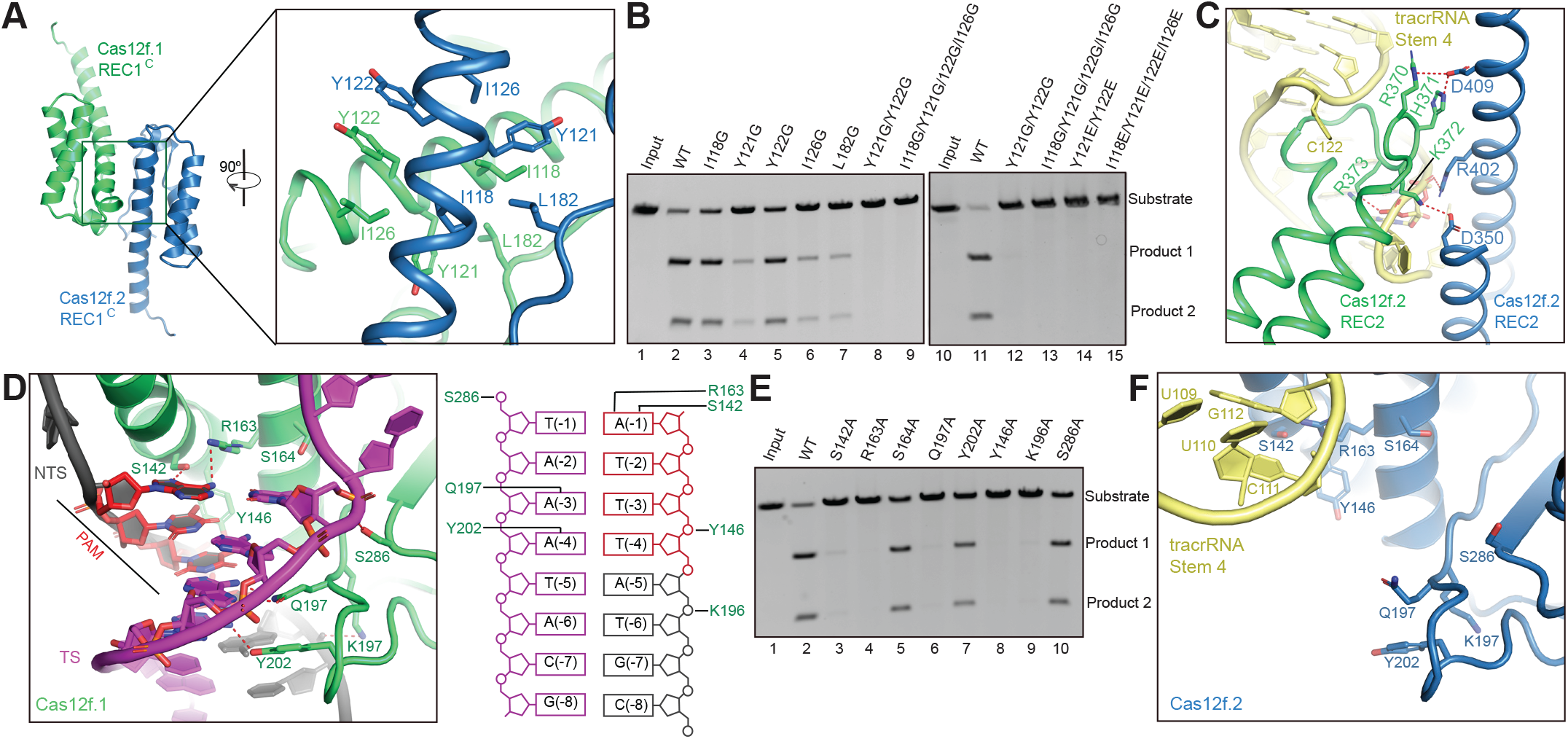
Dimerization, PAM recognition, and active site of Cas12f. **(A)** Dimerization interface of Cas12f mediated by REC1^C^. **(B)** Substrate DNA cleavage assay using wild-type Cas12f and Cas12f with mutations in the dimerization interface. The results shown are representative of three experiments. **(C)** interaction between two REC2 domains of Cas12f.1 and Cas12f.2. **(D)** Interactions between the PAM duplex and Cas12f.1. Schematic of the interactions is shown on the right. **(E)** Substrate DNA cleavage assay using wild-type Cas12f and alanine substitutions for residues involved in PAM recognition. The results shown are representative of three experiments. **(F)** PAM binding site of Cas12f.2.

### PAM recognition

The PAM sequence is recognized at the interface of REC1^C^ and the WED domain. The hydroxyl group of S142 and the guanidino group of R163 form two hydrogen bonds with base A (−1) of the TTTA PAM sequence in the non-target strand **(Fig. 3D)**. The amide group of Q197 forms a pair of hydrogen bonds with A (−3) of the target strand while Y202 forms a hydrogen bond with A (−4) of the target strand **(Fig. 3D)**. Alanine substitution of any of the residues reduces substrate cleavage activity with S142A, R163A, and Q197A almost completely abolishing activity **(Fig. 3E)**. In addition to the sequence-specific interactions, S286, Y146, and K196 also establish non-sequence-specific interactions with the PAM duplex **(Fig. 3D)**. Y146A and K196A mutations also severely reduce the complex’s ability to degrade substrate DNA **(Fig. 3E)**.

Notably, those residues are exposed to solvent in the other subunit, Cas12f.2, of the ternary complex; therefore, alanine substitutions of any of them should not impact substrate recognition **(Fig. 3F)**. Interestingly, Stem 4 of the sgRNA is located near the PAM recognition site in Cas12f.2, likely preventing substrate binding at this site **(Fig. 3F)**.

### crRNA-DNA heteroduplex recognition

The 19-20 bp crRNA-DNA heteroduplex is located in the central channel formed by Cas12f, similar to other Cas12 proteins **(Fig. S3)**. The heteroduplex is recognized by positively charged residues from both Cas12f.1 and Cas12f.2 while the non-target strand is held predominantly by the N-terminal half of Cas12f.2 **(Figs. 1A and S4A,B)**. The PAM proximal end of the heteroduplex is primarily recognized by Cas12f.1, whereas the PAM distal end is bound to Cas12f.2 **(Fig. S4A,B)**. The PAM distal end is capped by F341 and R343 from the RuvC domain of Cas12f.2 **(Fig. S4A,B)**. Single alanine substitutions for the residues involved in the recognition of the crRNA-DNA heteroduplex mostly result in modest reductions in the cleavage activity of Cas12f **(Fig. S4C)**. However, alanine substitution of R396 severely reduces the substrate cleavage activity **(Fig. S4C)**. R396 engages the phosphate group of position +8 of the target DNA strand, a position shown to be a critical checkpoint for Cas12a ^19,37^.

### Nuclease site of Cas12f

The conserved triplet of acidic residues (D326, E422, and D510) from the RuvC domain is located in the interface between the RuvC and Nuc domains **(Fig. 4A)**. Located in the active site is also R490 from the Nuc domain, and alanine substitution of this residue results in loss of cleavage activity ^8^. Lying on top of the acidic residues is the lid motif, which plays a vital role in regulating the RuvC active site ^25^. Interestingly, the lid motif in Cas12f.1 is in an open conformation, in correspondence to the crRNA-target DNA heteroduplex formation **(Fig. 4A)**. However, the lid motif in Cas12f.2 is in a closed conformation, although this active site is closer to the 5’ end of the target strand **(Fig. 4B)**. Two purine bases from the Stem 2 of tracrRNA, G(24) and A(62), insert into the inactive RuvC catalytic pocket, likely further inhibiting substrate access **(Figs. 4B and S4D)**. This observation indicates that only the RuvC domain in Cas12f.1 is activated upon target DNA binding.

**Figure 4.**
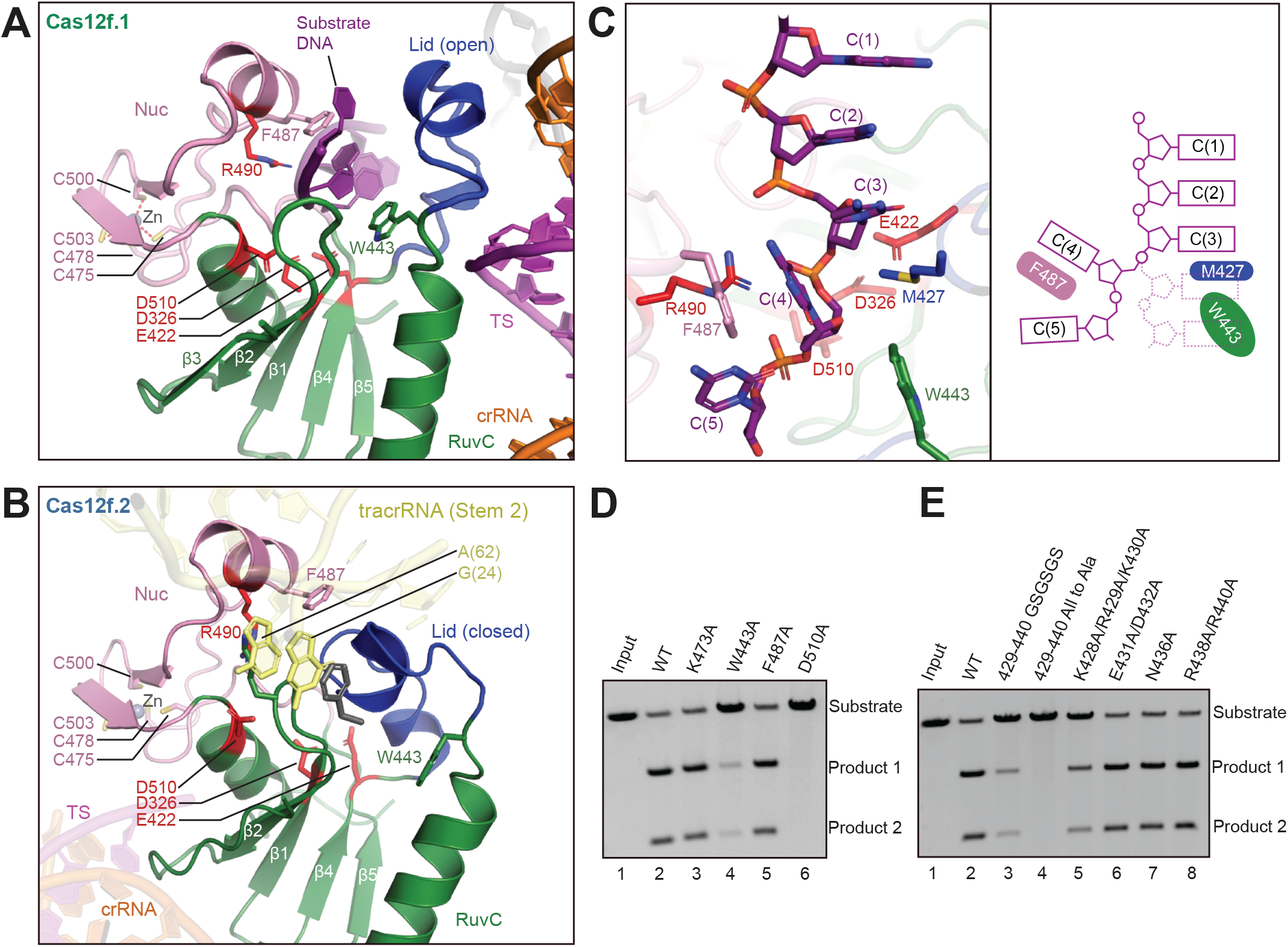
Nuclease site of Cas12f. **(A)** The nuclease site of the RuvC domain of Cas12f.1 in an open conformation. The acidic residues (D326, E422, and D510) from the RuvC domain and R490 from the Nuc domain are colored in red while the lid motif is in blue. Poly-C substrate is in purple. **(B)** The nuclease site of the RuvC domain of Cas12f.2 in a closed conformation. **(C)** Substrate DNA trapped in the nuclease site of Cas12f.1. Schematic is shown on the right. **(D)** Substrate DNA cleavage assay using wild-type Cas12f and various mutations around the substrate. The results shown are representative of three experiments. **(E)** Substrate DNA cleavage assay using wild-type Cas12f and various mutations in the lid motif. The results shown are representative of three experiments.

Interestingly, we observed an extended density assigned as the substrate DNA trapped in the RuvC active site of Cas12f.1, likely from excess DNA oligos used in complex assembly. The resolution does not allow for unambiguous assignment of bases but was clear enough for us to build a 5-nt poly-C model **(Fig. 4C)**. The backbone of the substrate is located in proximity to the triplet of acidic residues with R490 from the Nuc domain sitting on the other side of the backbone **(Fig. 4A)**. The stacking of bases in the substrate is broken between C(3) and C(4) due to the side chains of M427 and W433 occupying the space of base C(4). Consequently, base C(4) rotates by ∼ 90° and packs against the side chain of F487 **(Fig. 4C)**. The rotation of C(4) and the close proximity to the triplet of acidic residues indicate the phosphate group connecting C(3) and C(4) is the scissile phosphate targeted for cleavage. This configuration of the substrate DNA in the RuvC active site is consistent with previous observations in Cas9 ^38^, Cas12b ^21^ and Cas12i ^25^. Alanine substitution of W443 significantly reduces cleavage activity of Cas12f while F487 shows minor effect, suggesting that W443 plays a dominant role in positioning the substrate DNA for cleavage **(Fig. 4D)**.

The lid motif bridges the substrate and the crRNA-target DNA heteroduplex. Replacement of the lid motif with alanine or a GSGSGS linker deactivates Cas12f **(Fig. 4E)**. These results add to our mechanistic understanding of substrate configuration and cleavage within the RuvC nuclease domain.

### Activation mechanism of Cas12f

To understand the mechanism of Cas12f activation by target DNA, we determined the cryo-EM structure of Cas12f-sgRNA binary complex at 3.9 Å **(Figs. 5A, S5, and Table 1)**. This structure reveals a 5-nt pre-ordered seed sequence in the crRNA adjacent to the PAM duplex **(Fig. S5J)**. The 5’ end seed sequence was also observed in other Cas12 nucleases, including Cas12a ^16^, Cas12b ^21,22^, and Cas12i ^25^. The binary complex structure also allows us to reveal the conformational changes in Cas12f upon target DNA recognition **(Fig. 5B,C, and Movie S1)**. The most significant conformational changes happen in the C-terminal half of Cas12f.1 (RuvC, REC2, and Nuc) and N-terminal half of Cas12f.2 (REC1 and WED); both move outward upon the formation of the crRNA-target DNA heteroduplex between them. The lid motif of Cas12f.1 transits from the closed to open conformation, exposing the active site to accommodate ssDNA substrate. However, almost no conformational changes are observed in the RuvC domain of Cas12f.2, further suggesting that Cas12f.2 is not responsible for substrate cleavage. The conformational changes, particularly in the lid motif of Cas12f.1, are similar to those observed in Cas12a, Cas12b, and Cas12i ^25^. Although both copies of the Cas12f effector protein are necessary for the complex’s functionality, this evidence suggests that Cas12f still adopts a conserved mechanism for activation of the RuvC nuclease site like other type V nucleases.

**Figure 5.**
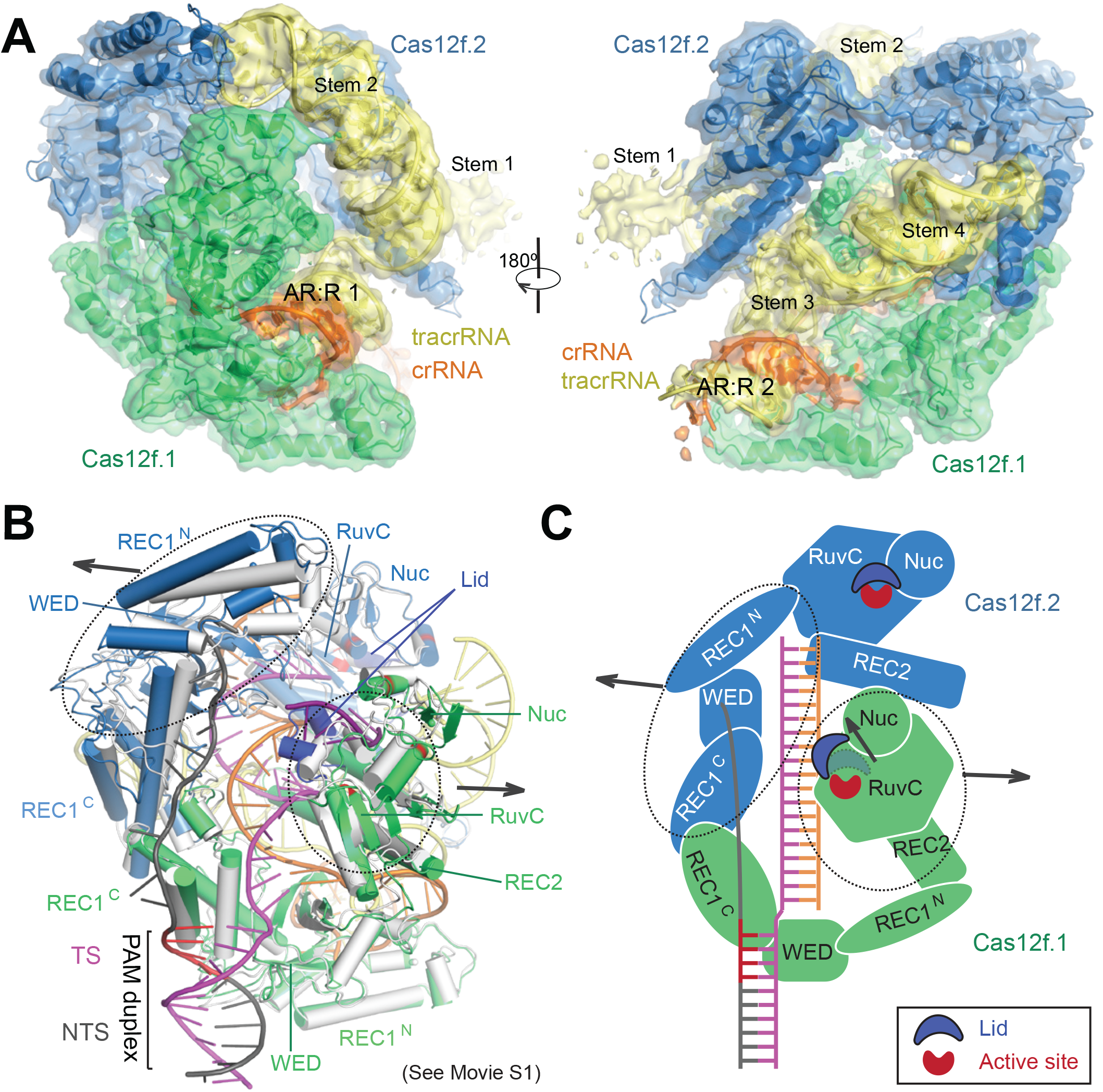
Structure of the Cas12f-sgRNA complex. **(A)** Cryo-EM map of the Cas12f-sgRNA complex at 3.9 Å in two views with each subunit color coded as in **Fig. 1A**. The map is shown in transparent surface while the underlying atomic model is shown in cartoon. **(B)** Structural superimposition of Cas12f-sgRNA (grey) and Cas12f-sgRNA-target DNA (colored) complexes. Conformational changes in Cas12f domains and sgRNA after recognition of target DNA are indicated by arrows. **(C)** Schematic showing the conformational changes in Cas12f upon target DNA binding.

## DISCUSSION

In this paper, we show that two copies of Cas12f bind to one sgRNA for target recognition and cleavage. Dimerization of Cas12f is likely to compensate for the small size of Cas12f, allowing for recognition of the ∼ 20-bp crRNA-target DNA duplex, which is a conserved length for substrate recognition in most class 2 CRISPR-Cas systems **(Fig. S3)**.

The most notable differences between Cas12f and other type V effectors are the lengths of the REC1 and REC2 domains. The REC1 domain of Cas12f is composed of ∼170 a.a., in comparison to ∼300 a.a. in other Cas12 proteins (315 aa in Cas12a ^18^, ∼377 a.a. in Cas12b ^21^, ∼276 a.a. in Cas12e ^24^, and ∼353 a.a. in Cas12i ^25^). Additionally, the REC2 domain in Cas12f is composed of ∼68 a.a., in comparison to ∼200 a.a. in other Cas12 proteins (∼252 a.a. in Cas12a ^18^, ∼200 a.a. in Cas12b ^21^, ∼177 a.a. in Cas12e ^24^, and ∼203 a.a. in Cas12i ^25^). Both the REC1 and REC2 domains are involved in the recognition and stabilization of the 20-bp crRNA-target DNA duplex, formation of which induces conformational changes required for activation of the RuvC domain. Minimal lengths of the REC1 and REC2 domains are thought to be indispensable for their proper function. Despite the miniature size of its REC1 and REC2 domains, dimerization renders Cas12f an effective RNA-guided nuclease similar to other Cas12 proteins. In detail, Cas12f.1 functions as a conventional Cas12 effector which contributes the canonical RuvC, Nuc, and WED domains. The combination of the REC1 domain of Cas12f.1 and the REC1 and WED domains of Cas12f.2 is structurally and functionally equivalent to the REC1 domains of other Cas12 proteins while the combination of the REC2 domain of Cas12f.1 and the RuvC, REC2 and Nuc domains of Cas12f.2 is a structurally and functionally equivalent to the REC2 domain of other Cas12 proteins. Consistent with this idea, F341 from the RuvC domain of Cas12f.2 packs against G(20) of the crRNA **(Fig. S4A,B)**, likely regulating the length of the crRNA-target DNA duplex similar to W382 in the REC2 domain of the *Acidaminococcus sp*. Cas12a ^39^.

Although two Cas12f molecules are required for target recognition and cleavage, Cas12f adopts the conserved activation mechanism of the type V nucleases that requires coordinated conformational changes induced by the formation of the crRNA-target DNA heteroduplex. In summary, our results unravel the mechanism of Cas12f and add to our understanding of mechanisms behind the diverse type V CRISPR-Cas effectors.

## ACKNOWLEDGMENTS

We thank Thomas Klose for help with cryo-EM, Steven Wilson for computation, and Clinton Gabel for helpful comments. This work made use of the Purdue Cryo-EM Facility. This work was supported by the NIH grant R01GM138675 to L.C.

## AUTHOR CONTRIBUTIONS

L.C. supervised the study. R.X. and Z.L. prepared samples. Z.L., R.X., and L.C. collected and processed cryo-EM data. R.X., S.W, R.H., and Z.L. performed biochemical analysis. All authors analyzed the data. R.X., Z.L., and L.C prepared the manuscript with input from other authors.

## DATA AVAILABILITY

Cryo-EM reconstructions of Cas12f-sgRNA-target DNA and Cas12f-sgRNA complexes have been deposited in the Electron Microscopy Data Bank under the accession numbers EMD-23158 and EMD-23157, respectively. Coordinates for atomic models of Cas12f-sgRNA-target DNA and Cas12f-sgRNA complexes have been deposited in the Protein Data Bank under the accession numbers 7L49 and 7L48, respectively.

## References

1 Sorek, R., Lawrence, C. M. & Wiedenheft, B. CRISPR-mediated adaptive immune systems in bacteria and archaea. Annual review of biochemistry 82, 237–266 (2013).

2 Marraffini, L. A. CRISPR-Cas immunity in prokaryotes. Nature 526, 55 (2015).

3 Jiang, F. & Doudna, J. A. CRISPR-Cas9 Structures and Mechanisms. Annu Rev Biophys 46, 505–529, doi:10.1146/annurev-biophys-062215-010822 (2017).

4 Hille, F. et al. The Biology of CRISPR-Cas: Backward and Forward. Cell 172, 1239–1259, doi:10.1016/j.cell.2017.11.032 (2018).

5 Makarova, K. S. et al. Evolutionary classification of CRISPR-Cas systems: a burst of class 2 and derived variants. Nat Rev Microbiol 18, 67–83, doi:10.1038/s41579-019-0299-x (2020).

6 Yan, W. X. et al. Functionally diverse type V CRISPR-Cas systems. Science 363, 88–91, doi:10.1126/science.aav7271 (2019).

7 Strecker, J. et al. RNA-guided DNA insertion with CRISPR-associated transposases. Science 365, 48–53, doi:10.1126/science.aax9181 (2019).

8 Harrington, L. B. et al. Programmed DNA destruction by miniature CRISPR-Cas14 enzymes. Science 362, 839–842, doi:10.1126/science.aav4294 (2018).

9 Karvelis, T. et al. PAM recognition by miniature CRISPR-Cas12f nucleases triggers programmable double-stranded DNA target cleavage. Nucleic Acids Res 48, 5016–5023, doi:10.1093/nar/gkaa208 (2020).

10 Dong, D. et al. The crystal structure of Cpf1 in complex with CRISPR RNA. Nature 532, 522 (2016).

11 Gao, P., Yang, H., Rajashankar, K. R., Huang, Z. & Patel, D. J. Type V CRISPR-Cas Cpf1 endonuclease employs a unique mechanism for crRNA-mediated target DNA recognition. Cell research 26, 901 (2016).

12 Yamano, T. et al. Crystal structure of Cpf1 in complex with guide RNA and target DNA. Cell 165, 949–962 (2016).

13 Yamano, T. et al. Structural basis for the canonical and non-canonical PAM recognition by CRISPR-Cpf1. Molecular cell 67, 633-645. e633 (2017).

14 Stella, S., Alcón, P. & Montoya, G. Structure of the Cpf1 endonuclease R-loop complex after target DNA cleavage. Nature 546, 559 (2017).

15 Stella, S. et al. Conformational Activation Promotes CRISPR-Cas12a Catalysis and Resetting of the Endonuclease Activity. Cell (2018).

16 Swarts, D. C., van der Oost, J. & Jinek, M. Structural Basis for Guide RNA Processing and Seed-Dependent DNA Targeting by CRISPR-Cas12a. Mol Cell 66, 221–233 e224, doi:10.1016/j.molcel.2017.03.016 (2017).

17 Nishimasu, H. et al. Structural basis for the altered PAM recognition by engineered CRISPR-Cpf1. Molecular cell 67, 139-147. e132 (2017).

18 Swarts, D. C. & Jinek, M. Mechanistic Insights into the cis- and trans-Acting DNase Activities of Cas12a. Mol Cell 73, 589–600 e584, doi:10.1016/j.molcel.2018.11.021 (2019).

19 Zhang, H. et al. Structural Basis for the Inhibition of CRISPR-Cas12a by Anti-CRISPR Proteins. Cell Host Microbe 25, 815–826 e814, doi:10.1016/j.chom.2019.05.004 (2019).

20 Yamano, T. et al. Structural Basis for the Canonical and Non-canonical PAM Recognition by CRISPR-Cpf1. Mol Cell 67, 633–645 e633, doi:10.1016/j.molcel.2017.06.035 (2017).

21 Yang, H., Gao, P., Rajashankar, K. R. & Patel, D. J. PAM-Dependent Target DNA Recognition and Cleavage by C2c1 CRISPR-Cas Endonuclease. Cell 167, 1814–1828 e1812, doi:10.1016/j.cell.2016.11.053 (2016).

22 Liu, L. et al. C2c1-sgRNA Complex Structure Reveals RNA-Guided DNA Cleavage Mechanism. Mol Cell 65, 310–322, doi:10.1016/j.molcel.2016.11.040 (2017).

23 Wu, D., Guan, X., Zhu, Y., Ren, K. & Huang, Z. Structural basis of stringent PAM recognition by CRISPR-C2c1 in complex with sgRNA. Cell Res 27, 705–708, doi:10.1038/cr.2017.46 (2017).

24 Liu, J. J. et al. CasX enzymes comprise a distinct family of RNA-guided genome editors. Nature 566, 218–223, doi:10.1038/s41586-019-0908-x (2019).

25 Zhang, H., Li, Z., Xiao, R. & Chang, L. Mechanisms for target recognition and cleavage by the Cas12i RNA-guided endonuclease. Nat Struct Mol Biol, doi:10.1038/s41594-020-0499-0 (2020).

26 Suloway, C. et al. Automated molecular microscopy: the new Leginon system. J Struct Biol 151, 41–60, doi:10.1016/j.jsb.2005.03.010 (2005).

27 Zivanov, J. et al. New tools for automated high-resolution cryo-EM structure determination in RELION-3. Elife 7, doi:10.7554/eLife.42166 (2018).

28 Zheng, S. Q. et al. MotionCor2: anisotropic correction of beam-induced motion for improved cryo-electron microscopy. Nat Methods 14, 331–332, doi:10.1038/nmeth.4193 (2017).

29 Zhang, K. Gctf: Real-time CTF determination and correction. J Struct Biol 193, 1–12, doi:10.1016/j.jsb.2015.11.003 (2016).

30 Punjani, A., Rubinstein, J. L., Fleet, D. J. & Brubaker, M. A. cryoSPARC: algorithms for rapid unsupervised cryo-EM structure determination. Nat Methods 14, 290–296, doi:10.1038/nmeth.4169 (2017).

31 Emsley, P., Lohkamp, B., Scott, W. G. & Cowtan, K. Features and development of Coot. Acta Crystallogr D Biol Crystallogr 66, 486–501, doi:10.1107/S0907444910007493 (2010).

32 Jones, D. T. Protein secondary structure prediction based on position-specific scoring matrices. J Mol Biol 292, 195–202, doi:10.1006/jmbi.1999.3091 (1999).

33 Afonine, P. V. et al. Real-space refinement in PHENIX for cryo-EM and crystallography. Acta Crystallogr D Struct Biol 74, 531–544, doi:10.1107/S2059798318006551 (2018).

34 Tan, Y. Z. et al. Addressing preferred specimen orientation in single-particle cryo-EM through tilting. Nat Methods 14, 793–796, doi:10.1038/nmeth.4347 (2017).

35 Pettersen, E. F. et al. UCSF Chimera--a visualization system for exploratory research and analysis. J Comput Chem 25, 1605–1612, doi:10.1002/jcc.20084 (2004).

36 Takeda, S. N. et al. Structure of the miniature type V-F CRISPR-Cas effector enzyme. Mol Cell, doi:10.1016/j.molcel.2020.11.035 (2020).

37 Stella, S. et al. Conformational Activation Promotes CRISPR-Cas12a Catalysis and Resetting of the Endonuclease Activity. Cell 175, 1856–1871 e1821, doi:10.1016/j.cell.2018.10.045 (2018).

38 Jiang, F. et al. Structures of a CRISPR-Cas9 R-loop complex primed for DNA cleavage. Science 351, 867–871, doi:10.1126/science.aad8282 (2016).

39 Gao, P., Yang, H., Rajashankar, K. R., Huang, Z. & Patel, D. J. Type V CRISPR-Cas Cpf1 endonuclease employs a unique mechanism for crRNA-mediated target DNA recognition. Cell Res 26, 901–913, doi:10.1038/cr.2016.88 (2016).

