## Supplementary Materials for "Structural basis for the dimerization-dependent CRISPR-Cas12f nuclease"

### SUPPLEMENTARY FIGURES

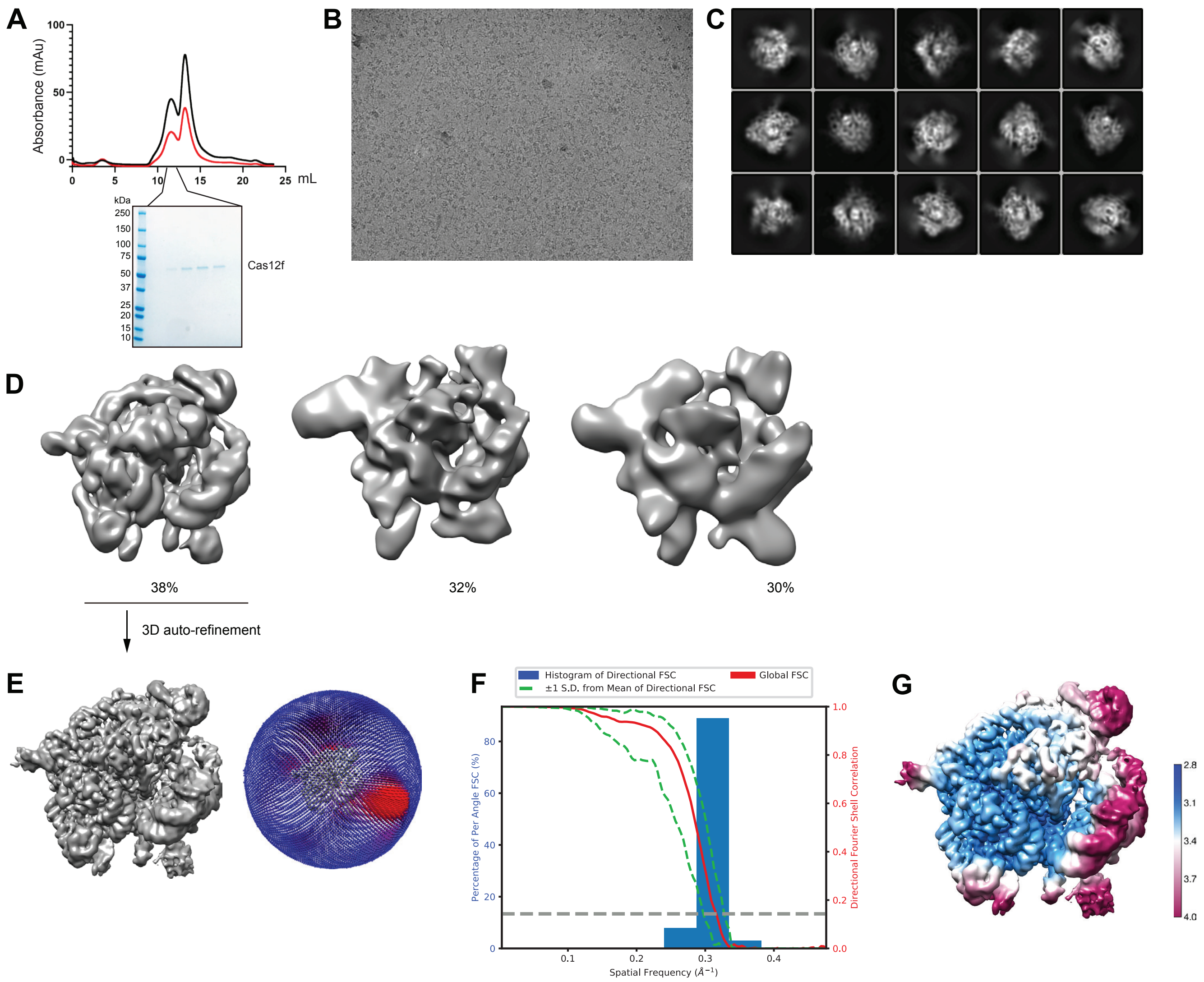


**Figure S1. Sample preparation and cryo-EM of the Cas12f-sgRNA-target DNA ternary complex.** **(A)** Purification of the Cas12f-sgRNA-target DNA ternary complex by size exclusion chromatography (SEC). UV absorbance curves at 280 nm and 260 nm are shown in red and black, respectively. SDS-PAGE analysis is shown below the SEC profile. **(B)** A representative raw cryo-EM micrograph of the Cas12f-sgRNA-target DNA complex from a total of 2450 micrographs. **(C)** Representative 2D class averages from a total of 100 images. **(D)** Three major classes from 3D classification. **(E)** 3D refinement for Class I with angular distribution of particles. **(F)** Plot of the global half-map FSC (solid red line) and spread of directional resolution values (±1σ from mean, green dotted lines; the blue bars indicate a histogram of 100 such values evenly sampled over the 3D FSC). FSC plot for the reconstruction suggests an average resolution of 3.1 Å. **(G)** Local resolution map for the reconstruction in **E**.

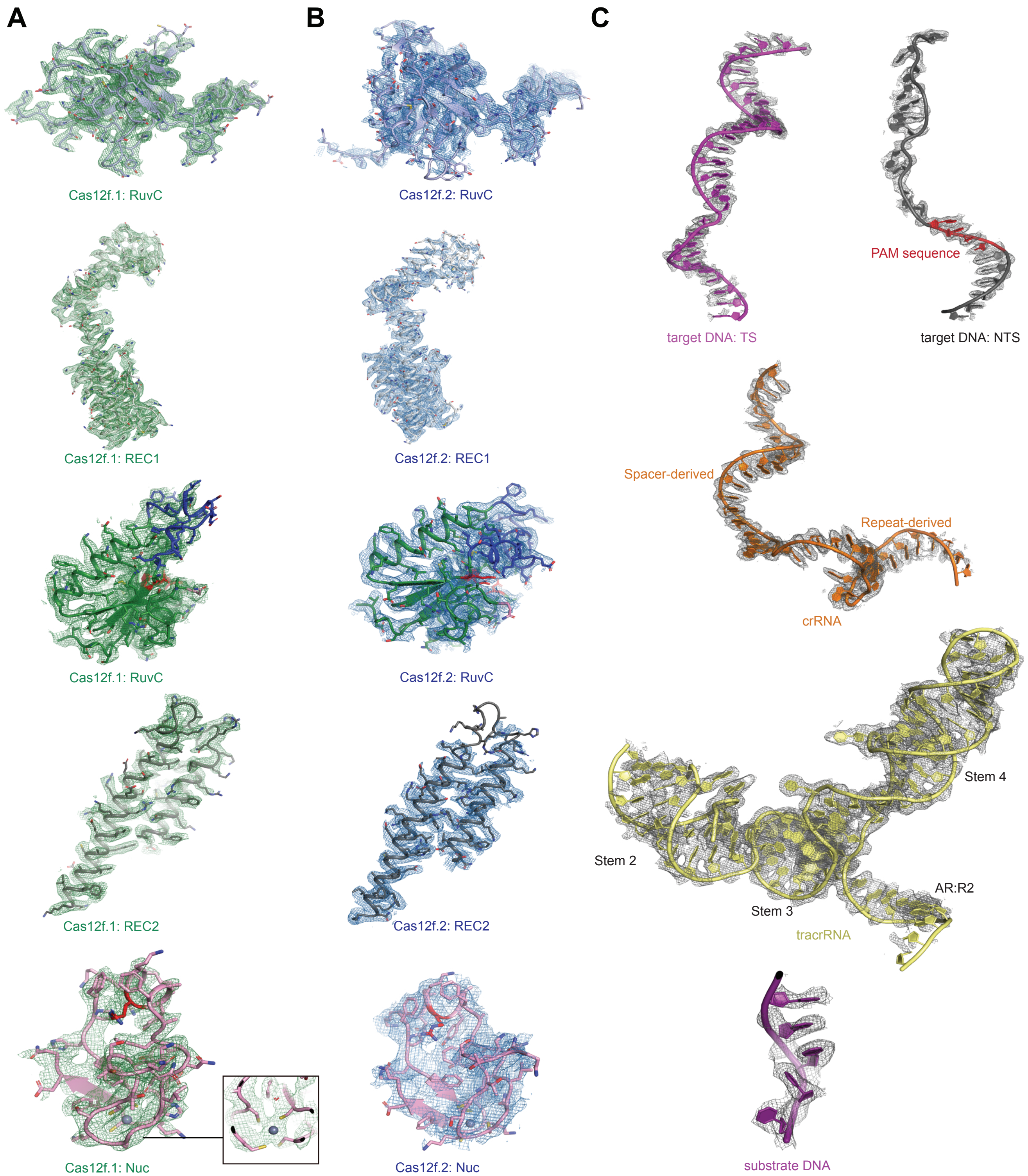


**Figure S2. Detailed cryo-EM density map of the Cas12f-sgRNA-target DNA complex with atomic model fitted in.** Cryo-EM density map (in mesh) of each domain in Cas12f.1 **(A)** and Cas12f.2 **(B)**, and each chain of nucleic acids **(C)** with atomic model fitted in.

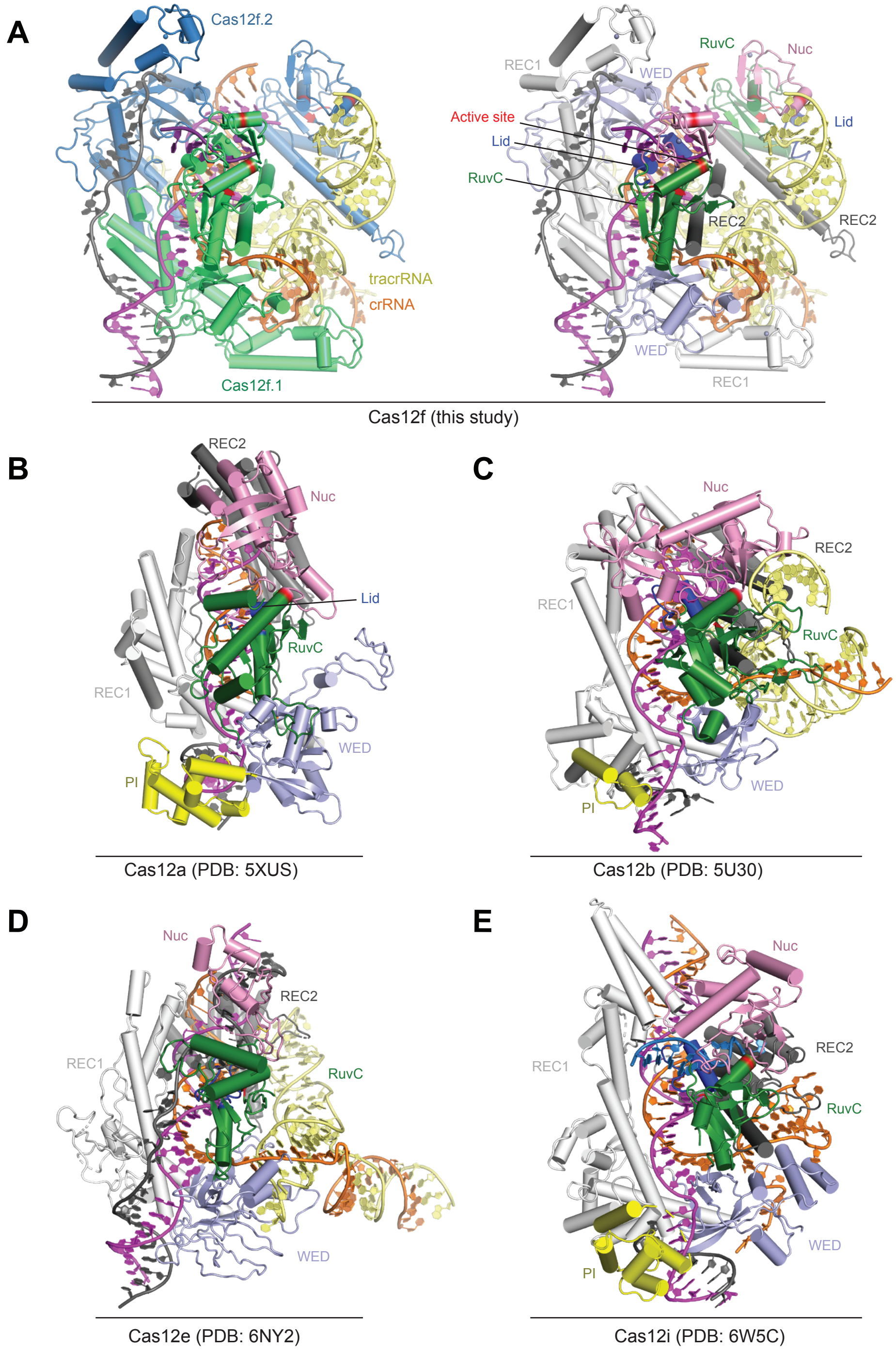


**Figure S3. Structural comparison between Cas12f and other Cas12 nucleases. (A)** Structure of Cas12f-sgRNA-target DNA complex. Left: two monomers of Cas12f are colored in green and blue, respectively. Right: each domain of Cas12f is color-coded as labeled. **(B-E)** Structure of Cas12a-crRNA-target DNA **(B)**, Cas12b-crRNA-target DNA **(C)**, Cas12e-crRNA-target DNA **(D)**, and Cas12i-crRNA-target DNA **(E)** with each domain color-coded as labeled.

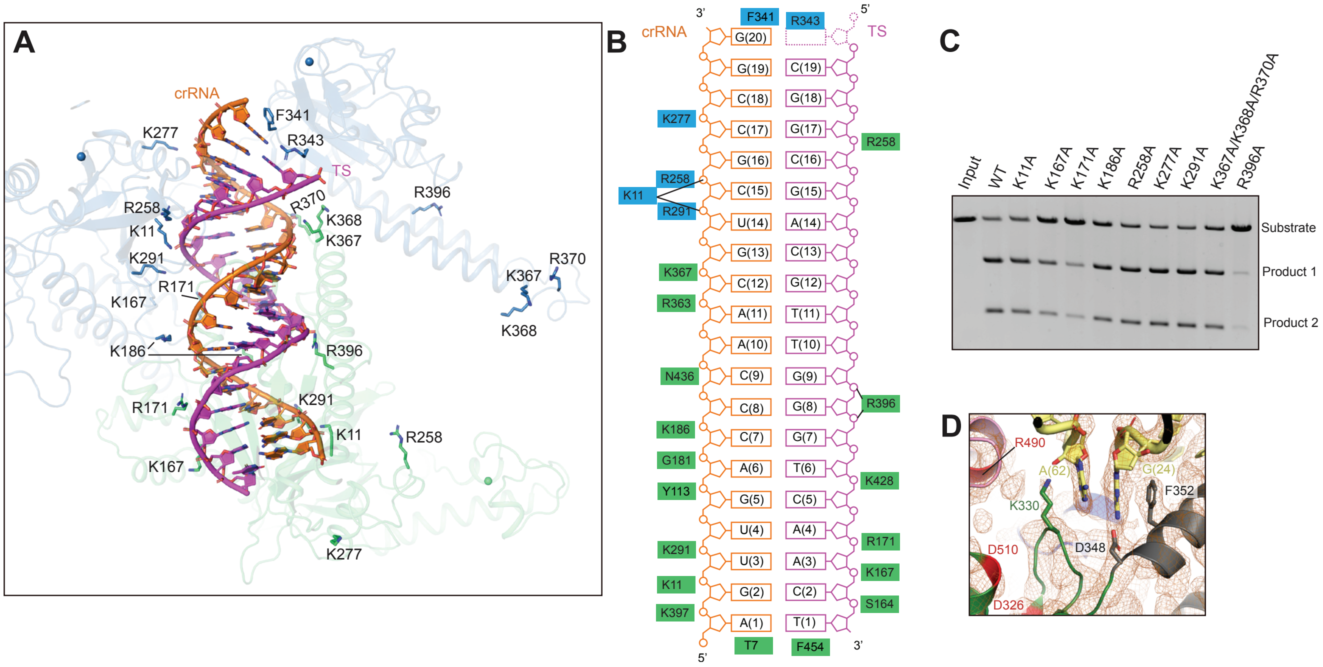


**Figure S4. Interactions between Cas12f proteins and nucleic acids in the Cas12f-sgRNA-target DNA complex. (A,B)** Structure **(A)** and schematic **(B)** showing the interactions between Cas12f and the crRNA-target DNA heteroduplex. **(C)** Substrate DNA cleavage assay using wild-type Cas12f and Cas12f with mutations in the positively charged residues which are involved in the recognition of the crRNA-target DNA heteroduplex. The results shown are representative of three experiments. **(D)** Interaction between tracrRNA and the RuvC active site of Cas12f.2.

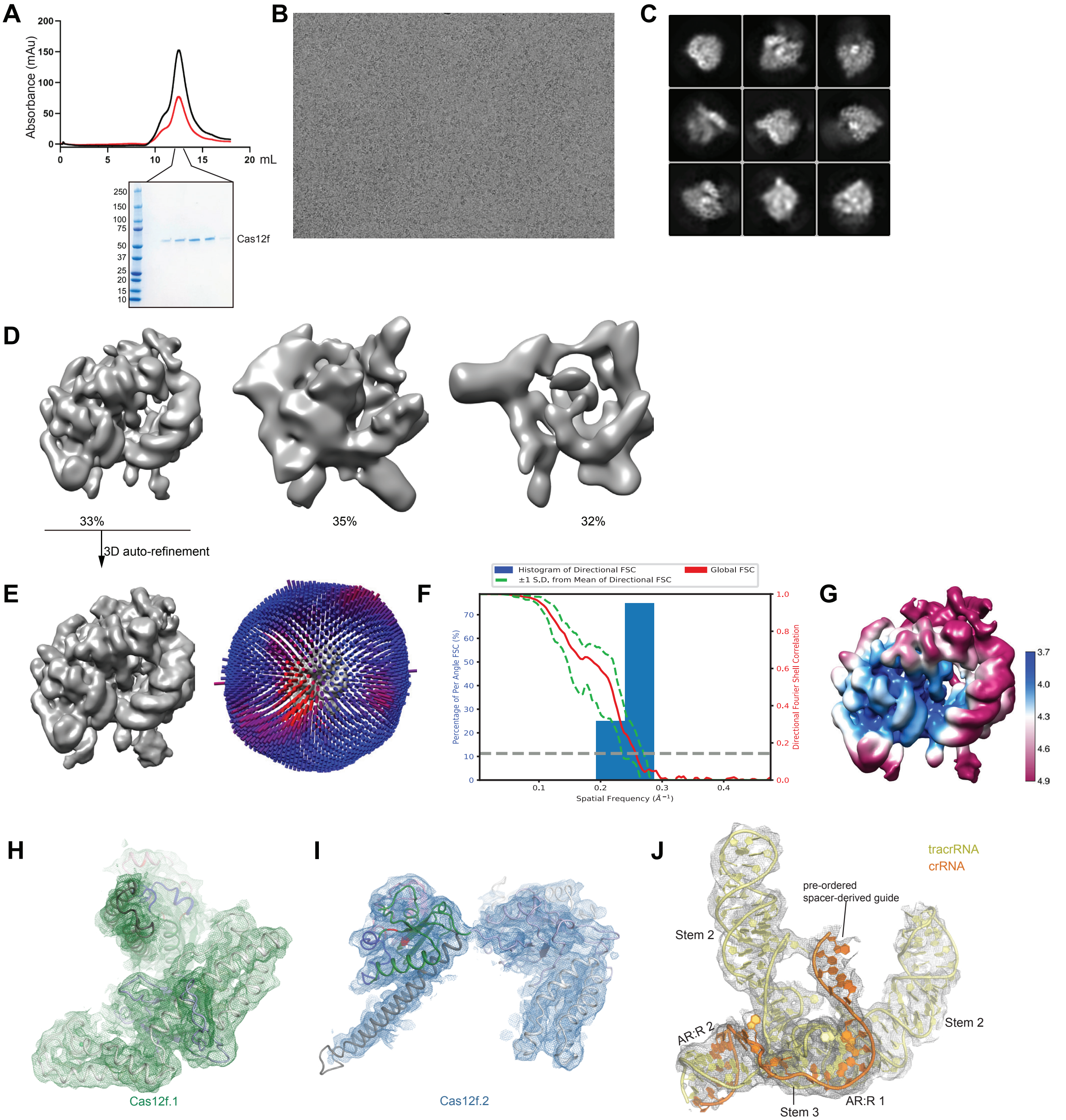


**Figure S5. Sample preparation and cryo-EM of the Cas12f-sgRNA binary complex. (A)** Purification of the Cas12f-sgRNA complex by size exclusion chromatography (SEC). UV absorbance curves at 280 nm and 260 nm are shown in red and black, respectively. SDS-PAGE analysis is shown below the SEC profile. **(B)** A representative raw cryo-EM micrograph of the Cas12f-sgRNA complex from a total of 1391 micrographs. **(C)** Representative 2D class averages from a total of 100 images. **(D)** Three major classes from 3D classification. **(E)** 3D refinement for Class I with angular distribution of particles. **(F)** Plot of the global half-map FSC (solid red line), and spread of directional resolution values (±1σ from mean, green dotted lines; the blue bars indicate a histogram of 100 such values evenly sampled over the 3D FSC). FSC plot for the reconstruction suggests an average resolution of 3.9 Å. **(G)** Local resolution map for the reconstruction in **E**. **(H-J)** Cryo-EM density map (in mesh) of Cas12f.1 **(H)**, Cas12f.2 **(I)**, and sgRNA **(J)** with atomic model fitted in.

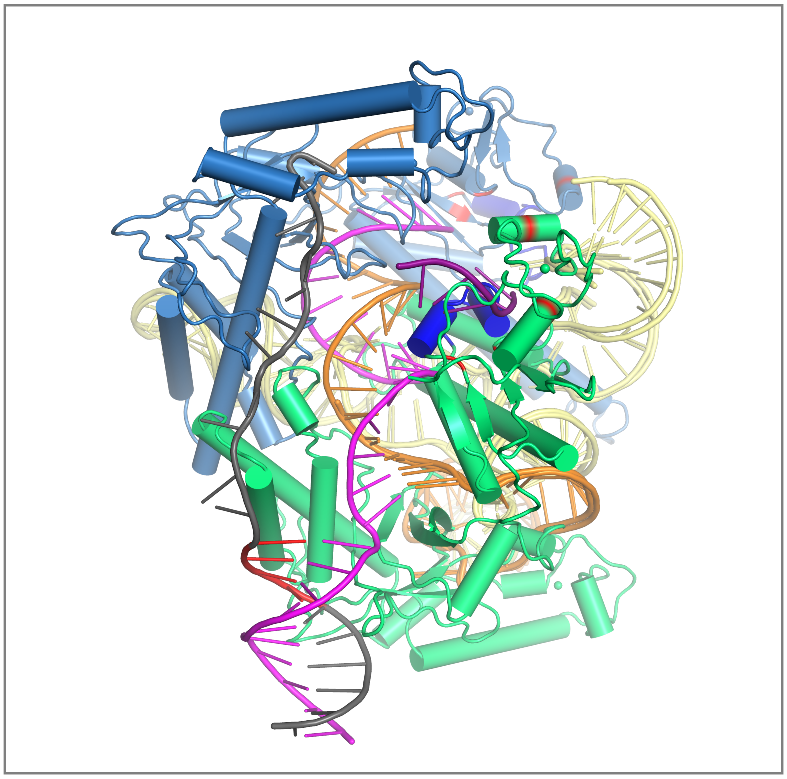

**Movie S1. Conformational changes in Cas12f-sgRNA upon target DNA binding.** This movie shows structural morphing of Cas12f-sgRNA before and after target DNA binding, in the same view as shown in **Fig. 5B**.

**Table S1. Oligonucleotides utilized in assembly Cas12f-sgRNA-target DNA complex**

| Oligonucleotides | Sequences |
|---|---|
| sgRNA | CUUCACUGAUAAAGUGGAGAACCGCUUCACCAAAAGCUGUCCCUUAGGGG AUUAGAACUUGAGUGAAGGUGGGCUGCUUGCAUCAGCCUAAUGUCGAGAA GUGCUUUCUUCGGAAAGUAACCCUCGAAACAAAUUCAUUUUUCCUCUCCAA UUCUGCACAAGAAAGUUGCAGAACCCGAAUAGACGAAUGAAGGAAUGCAAC AGUUGACCCAACGUCGCCGG |
| Target strand | GCTGATGCATCTAGATTGTGCACGCCGGCGACGTTGGGTCAACTTAAATACG TAAGGTGC |
| Non-target strand | GCACCTTACGTATTTAAGTTGACCCAACGTCGCCGGCGTGCACAATCTAGAT GCATCAGC |
